# A Multiplexed Fluorescent Microsphere Immunoassay of IgG Antibodies to the Individual Glycose Antigens of the *Neisseria* Lipooligosaccharide α Chain in Clients of an STI Clinic

**DOI:** 10.1101/2025.10.01.679747

**Authors:** Stephanie E. McLaughlin, Zhijie Yang, J. McLeod Griffiss

**Author notes:** **Corresponding Author**: Stephanie McLaughlin, MD MPH, Alternate Corresponding Author: J. McLeod Griffiss, MD, VAMC (111W1), 4150 Clement Street, San Francisco, CA 94121.

## Abstract

**Background:** To evaluate the immunogenicity of lipooligosaccharide (LOS) based gonococcal vaccines, we developed a multiplexed fluorescent microsphere immunoassay that distinguishes and quantifies antibodies that bind the various LOS antigens.

**Methods:** We coupled LOS from gonococcal strains 1291wt (nLc4 α chain), 1291a (Lc3 α chain) and 1291c (Lc2 α chain), to different fluorescent microsphere regions (BioRad) after disaggregation in 1% deoxycholate. The microspheres were incubated with sera from 37 participants in a study of gonococcal risk factors. Laser excited fluorescence from each region was converted to ng/mL of IgG with use of affinity purified human IgG specific for each LOS. IgG specific to the nLc4 terminal galactose was quantified by subtraction of Lc3 IgG from 1291wt IgG; that specific to the Lc3 terminal glucosamine by subtraction of Lc2 IgG from Lc3 IgG.

**Results:** Concentrations of nLc4, Lc3 and Lc2 IgG summed to the concentration of 1291wt IgG. Visual inspection of the data revealed a non-normally distributed, bimodal distribution of nLc4 IgG concentrations; 31/37 had a mean serum nLc4 IgG concentration of 6.22 ng/mL, and the other six had a mean nLc4 IgG concentration of 11.2 ng/ml. Upon statistical comparison of these groups, conducted prior to knowledge of infection status, those participants whose nLc4 IgG concentration centered around a higher mean, were more likely to be asymptomatic (*p* = 0.03) and were less likely to be infected (*p* = 0.05) than those with lower IgG concentrations.

**Conclusions:** This accurate LOS immunoassay can be expanded to include any number of LOS specificities.

**Short Summary:** We developed a multiplexed fluorescent microsphere immunoassay of IgG antibodies to glycose antigens of the *Neisseria* Lipooligosaccharide in clients of an STI clinic who were sexually exposed to gonorrhea.

## Background

During phase 1 of a gonococcal human challenge study conducted at Walter Reed Army Institute of Research[1], 15 men underwent intraurethral instillation of an ID_100_ of MS11mkC, a strain isolated from the male urethra during a previous human gonococcal challenge[2]. Nearly all, 14/15, developed gonorrhea. Of the 14 who were infected, ten delayed treatment for at least six days, per protocol, to allow for an neo-antibody response[3]. In phase 2 of the study, the 14 previously infected phase 1 subjects were re-challenged with an ID_50_ along with 10 naïve subjects. Five naïve subjects were infected, whereas only 2/10 of those who delayed treatment per protocol in phase 1 were re-infected[1]. Those who resisted re-infection were significantly more likely to have had a fourfold or greater increase in MS11mkC lipooligosaccharide (LOS) IgG than those who did not resist reinfection (p=0.036)[1]. Elevated LOS IgG persisted for the 12 weeks of follow-up[1]. This study demonstrated that LOS IgG induction by vaccination might well prevent GC in the male urethra, but the LOS antigen that induced the protective IgG was not sought in the study.

LOS are glycolipids in the outer membrane of bacteria that colonize mucosal surfaces of the genital and respiratory tracts[4]. LOS have been implicated in the pathogenesis of diseases caused by *Neisseria* and *Haemophilus*[2,5], and antibodies to them provide protection from disease[1]. LOS are triantennary glycolipids; the three glycose antennae arise from a Type II lipid A structure through a KDO linkage[5,6]. The longest antenna, termed the α chain[2], is comprised of 2-5 glycoses. *N. gonorrhoeae* in the polynuclear leukocytes (PMNs) within the urethral discharge of men with gonorrhea make LOS with lacto-*N*-neotetraose (Paragloboside)[7] α chains[8]. Protective antibodies bind α chain antigens[1,9].

LOS α chains are highly variable, as three of the five genes in the biosynthetic operon contain poly-G tracts that are subject to slip-strand mispairing[10]. Each bacterium makes LOS with different α chains[11]. Initially it was thought that each LOS molecule presented a single antigen centered on the non-reducing terminal glycose – the immunodominant glycose – as modified by various basal substitutions[12]. According to this notion, the diversity of LOS antibodies resulted from the diversity of LOS glycoforms, and individual antibodies could be detected and quantified by using different LOS molecules[13].

However, studies of gonococcal immunity found that LOS that had truncated α chains and elongated β chains presented multiple antigens that were recognized by human IgG and were bactericidal for these strains[14]. A study of meningococcal immunity that used a series of pyocin-selected mutants of gonococcal strain 1291 that have sequential deletions from its lacto-*N-* neotetraose (nLc4) α chain[15] (Figure 1) found that each glycose within the chain conformed a distinct antigen[9]. Thus, the diversity of LOS antibodies results from the diversity of antigens within each different α and β chain. Furthermore, antibodies that bind to the different glycose antigens within an α chain may have different functional activity[9]. Human IgG that binds the terminal Gal of nLc4 (nLc4 IgG) are not bactericidal for the most prevalent, L3,7, strains of *N. meningitidis* in the presence of human complement, whereas those that bind the sub-terminal GlcNAc (Lc3 IgG) are bactericidal for these strains[9]. This complexity complicates the central question: what is the specificity of the LOS IgG that protects humans from gonococcal infection[1]?

**Figure 1.**
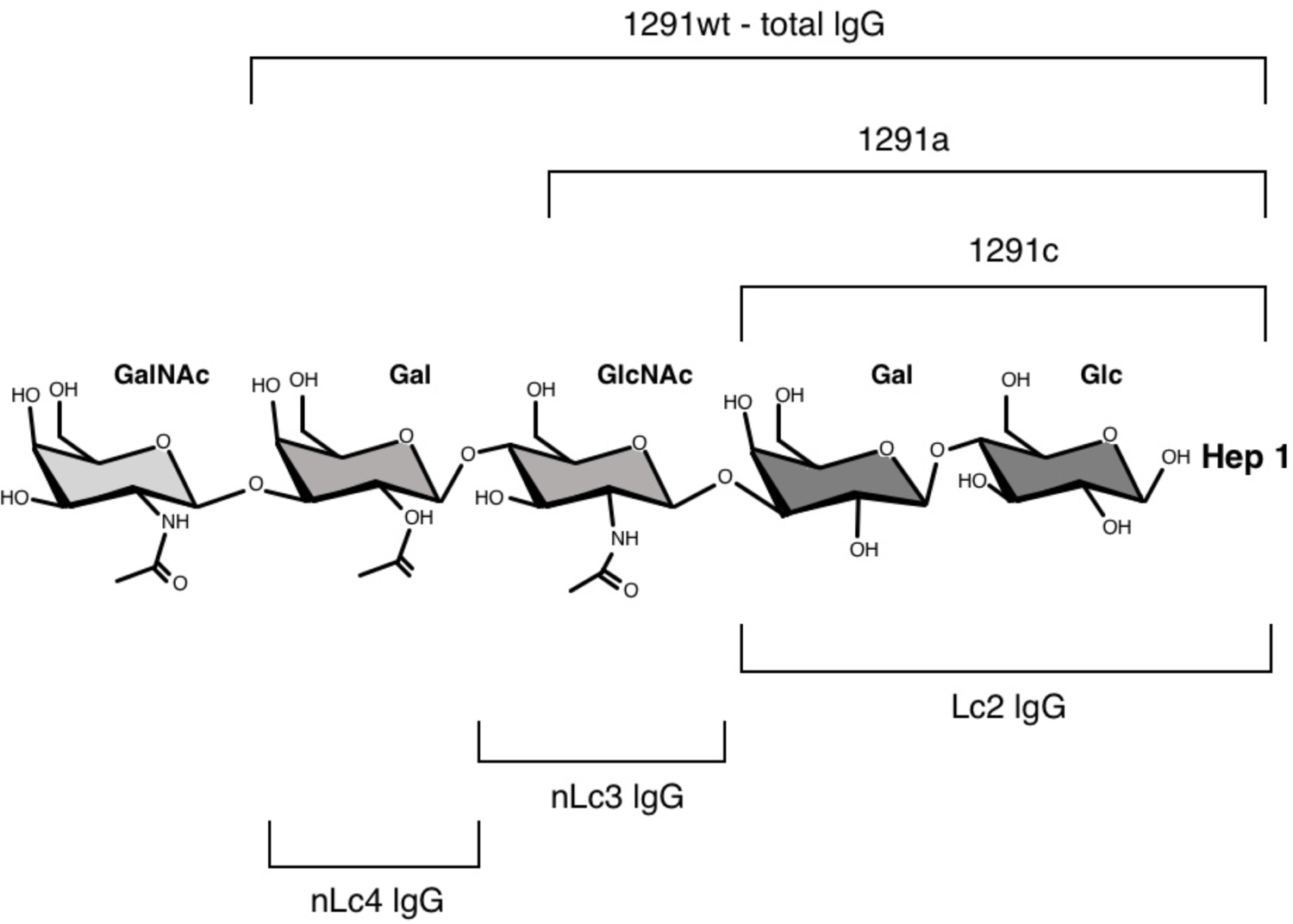
A schematic of the full-length gonococcal LOS α chain^1,2^(GalNAc-nLc4) that shows the specificities of the IgGs quantified in this study. *LgtD,* the gene that encodes a β-galactosaminyl transferase, is mostly OOF in the discharge of infected men. It shifts into frame during DNA and RNA replication. IgG specific for the terminal *N*-acetyl galactosamine (GalNAc) was not quantified in the study. 1291wt has a lacto-*N-*neotetraose (nLc4) α chain; 1291a and 1291c have sequential deletions of galactose (Gal) and *N*-acetyl glucosamine (GlcNAc), respectively. This results in, respectively, nLc3 and Lc2 α chains. 1291wt IgG consists of all the IgG specificities in a serum that bind any of the various antigens presented by the complete LOS molecule, including the basal glycoses and Lipid A; nLc4 IgG is specific for the nLc4 terminal Gal, Lc3 IgG is specific for the subterminal GlcNAc and Lc2 IgG for the internal β-lactose along with the other basal glycose and lipoidal antigens. ^1^The abbreviations used are: LOS, lipooligosaccharide(s); nLc4^3^, lacto-*N*-neotetraose; Lc3^3^, lacto-*N*-neotriaose; Lc2^3^, lactose; PEA, phosphoethanolamine; Gal, galactose; Glc, glucose; GlcNAc, *N*-acetyl glucosamine; GalNAc, *N*-acetyl galactosamine. ^2^Designations for LOS glycoses follow the recommendations of the International Union of Pure and Applied Chemistry/International Union of Biochemistry and Molecular Biology (IUPAC-IUB) Joint Commission on Biochemical Nomenclature, Nomenclature of Glycolipids (1997). These recommendations can be found at the IUPAC-IUB Web site.

To investigate the LOS immune response to the different nLc4 glycose antigens during gonococcal infection we developed a multiplexed fluorescent microsphere immunoassay. We used this assay to compare concentrations of nLc4, Lc3 and Lc2 IgG in sera of sexual contacts (GC-contacts) of index cisgender men (cismen) and cisgender women (ciswomen) with gonococcal infections who were enrolled in a study of risk factors for acquisition of infection during exposure[16]. We focused on nLc4 IgG and Lc3 IgGs[8,9].

## Methods

### Patient Consent Statement and Sample Storage

The study was approved by the Johns Hopkins University Institutional Review Board (NA-00004860), and written informed consent was obtained from all participants prior to study enrollment. Participants were compensated with gift cards of minimal value for their time. Participant sera were stored at −80C. *Multiplex assay* LOS purified (using the previously described hot phenol extraction method[17]) from the gonococcal 1291 mutants[15] that expressed nLc4 (1291wt), Lc3 (1291a) and Lc2 (1291c) LOS α chains[15] (Figure 1) were each coupled to 100 µL of Bio-Rad supplied fluorescent microspheres (Bio-Plex^®^ COOH Beads (Bio-Rad Laboratories, Inc, USA)). Coupling was achieved via carbodiimide reactions involving the primary amino groups of the LOS basal phosphoethanolamine (PEA) moieties and the carboxyl functional groups on the bead surfaces with use of the Amine Coupling Kit supplied by Bio-Rad. After coupling, beads were washed 2X with 500 µL of 1% deoxycholate to disaggregate micelles that contained LOS molecules that were not covalently bound[17], followed by washing in 500 µL of phosphate-buffered saline (PBS) to remove the deoxycholate. Each LOS was coupled to beads of a different fluorescence region[18]. Biotinylated monoclonal mouse anti-human IgG was the secondary antibody.

Two µL of each region microspheres were added to the different wells in the array, followed by 50 µL of LOS IgG standard, patient serum or blanks. Arrays were covered with aluminum foil, shaken at RT for two hours and washed with 100 µL of PBST after which 50 µL of a 1:1500 dilution of the secondary antibody was added and the array was shaken at RT for 30 minutes. Fifty µL of 1:500 dilution of phycoerythrin was then added to each well, and the array was again shaken at RT for 30 minutes. After washing 3X with 100 µL of PBST in each well, laser excited fluorescence from each region was recorded in a Bio-Plex^®^ 200 suspension array reader (Bio-Rad Laboratories, Inc, USA).

To convert the fluorescence recorded for each serum in the assay to IgG concentration, we included in each assay standard solutions containing known concentrations of affinity-purified human IgGs specific for each LOS α chain. We constructed curves that expressed the relationship between the IgG concentrations in each standard solution and the fluorescence readout of that solution’s binding to its LOS in the assay. The fluorescence read-outs for each patient’s serum were converted to IgG concentration by extrapolation from these standard curves.

The 1291 LOS region was assumed to have captured all the 1291 LOS IgG in that aliquot of the patient’s serum. We then quantified IgG that bound the nLc4 terminal β-Gal by subtracting the Lc3 IgG concentration from 1291 IgG concentration (nLc4 IgG). Similarly, we quantified IgG that bound the Lc3 terminal β-GlcNAc by subtracting Lc2 LOS IgG from Lc3 IgG (Lc3 IgG) (Figure 1).

We assumed that IgG concentrations in the sera of infected individuals that were seen within 6 days of exposure represented pre-exposure concentrations, whereas concentrations in sera drawn 7 or more days after exposure could contain IgG made in response to the infection[1].

### Standard Solutions for Multiplex Assay

We purified human IgG specific for 1291 LOS (nLc4 α chain), 1291a LOS (Lc3 α chain) and 1291c LOS (truncated basal α chain terminating in β-Lactose) (Figure 1) with a modification of the immuno-affinity columns described previously[9].

Briefly, LOS purified from 1291wt, 1291a and 1291c[15] were each coupled to epoxy-activated Sepharose 6B (Amersham Biosciences) through epoxy linkages to basal PEAs after disaggregation in 3% deoxycholate[17]. Coupling through the basal PEA moieties ensures that the conformation of the oligosacharide α chains in aqueous solutions is not affected.

We then purified human IgG specific for each LOS from commercial IVIG by mixing 500 µL of a 1:10 dilution of IVIG with 200 µL of each Sepharose affinity gel and allowing the IgG to bind the LOS during gentle rocking at RT for 10 m after which we spun the slurry at 0.1g for 1 m and let it sit undisturbed for a few minutes. We then carefully removed the supernatant until about 100 µL remained and washed the beads 3X with 1 mL PBS, removing the supernatant after each wash as above. After washing, the bound IgG was eluted with glycine-HCl (pH 2.3), and the pH of the eluate was immediately adjusted to pH 7–8 with 1 M Tris. The eluate was diluted 1:10 in Glycine-Tris (10:1) buffer.

To isolate IgG that bound all antigens within the 1291 LOS, IVIG was mixed with 1291 LOS-coupled Sepharose, and IgG, denominated 1291 IgG, was eluted from the beads. To isolate IgG that bound all the antigens presented by the Lc3 α chain, IVIG was mixed with the 1291a LOS-coupled Sepharose, and IgG, denominated 1291a IgG, was eluted. To isolate IgG that bound antigens within the truncated Lc2 α chain, IVIG was mixed with the 1291c LOS-coupled Sepharose, and IgG, denominated Lc2 IgG was eluted.

IgG in the eluates was quantified with use of the Easy-Titer Human IgG (H+L) Assay Kit with Pierce Human IgG Whole molecule ae standards (Thermo Fisher Scientific, Vacaville, CA), and the eluates were used as standards for quantifying IgG in patient sera.

To construct curves that expressed the relationships between IgG concentrations and fluorescence readout, we made serial 1:2 dilutions of each IgG preparation, starting at 5 µg/ml.

## Results

### Study Population

Thirty-seven GC-Contacts, enrolled at a Baltimore City Health Department (BCHD) Sexual Health Clinic from January, 2008 to May, 2012, contributed serum to this analysis; 46% were ciswomen (17/37) and 54% (20/37) cismen (see Table 1 for participant numbers of GC negative and positive cismen and ciswomen and see Supplemental Table 1 for LOS Ab concentrations and clinical and lab details by participant). The ages of GC-Contacts ranged from 18-39. The median age of the ciswomen and cismen was 23 and 27, respectively. All the cismen and 15 of the ciswomen were Black.

**Table 1.**
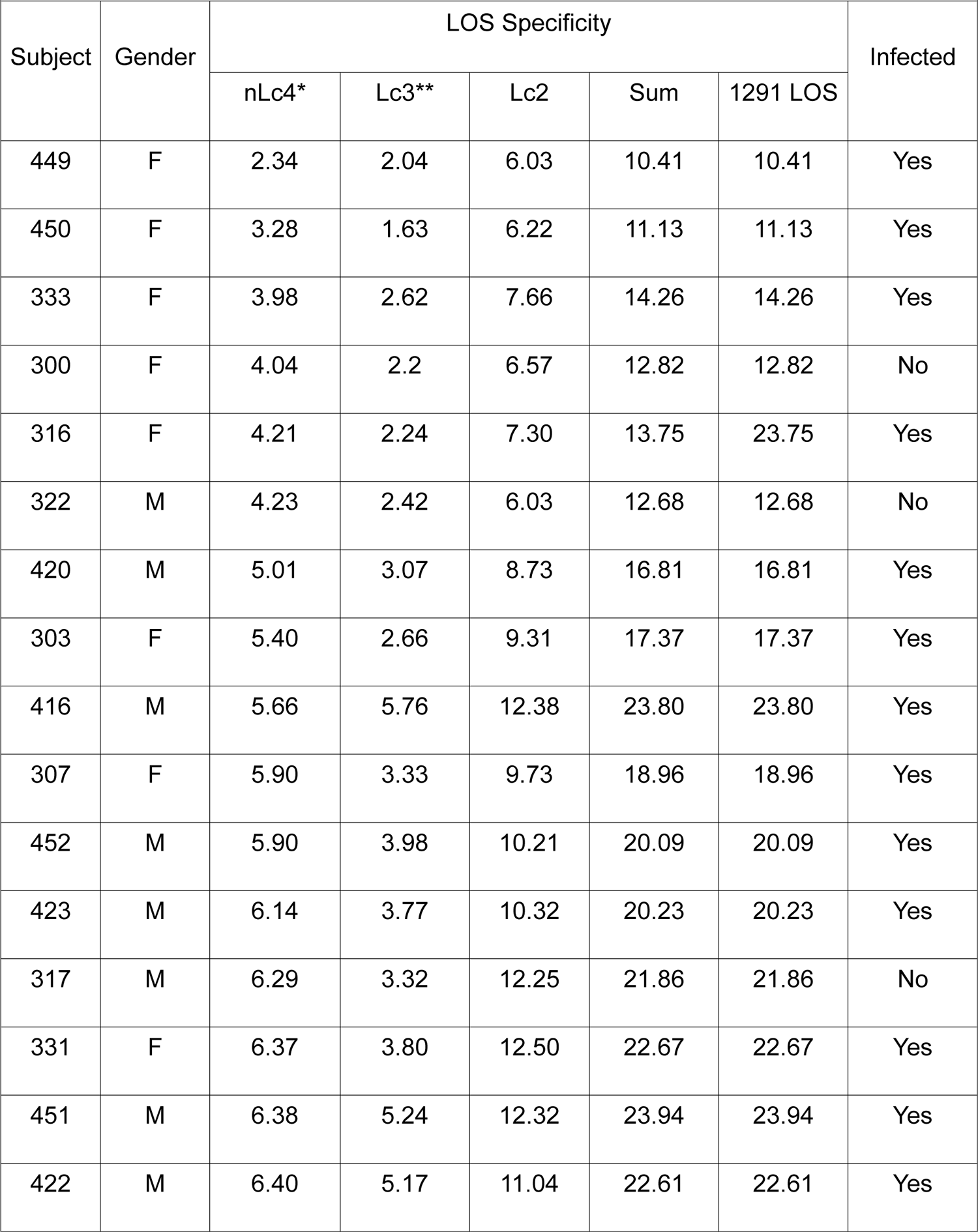
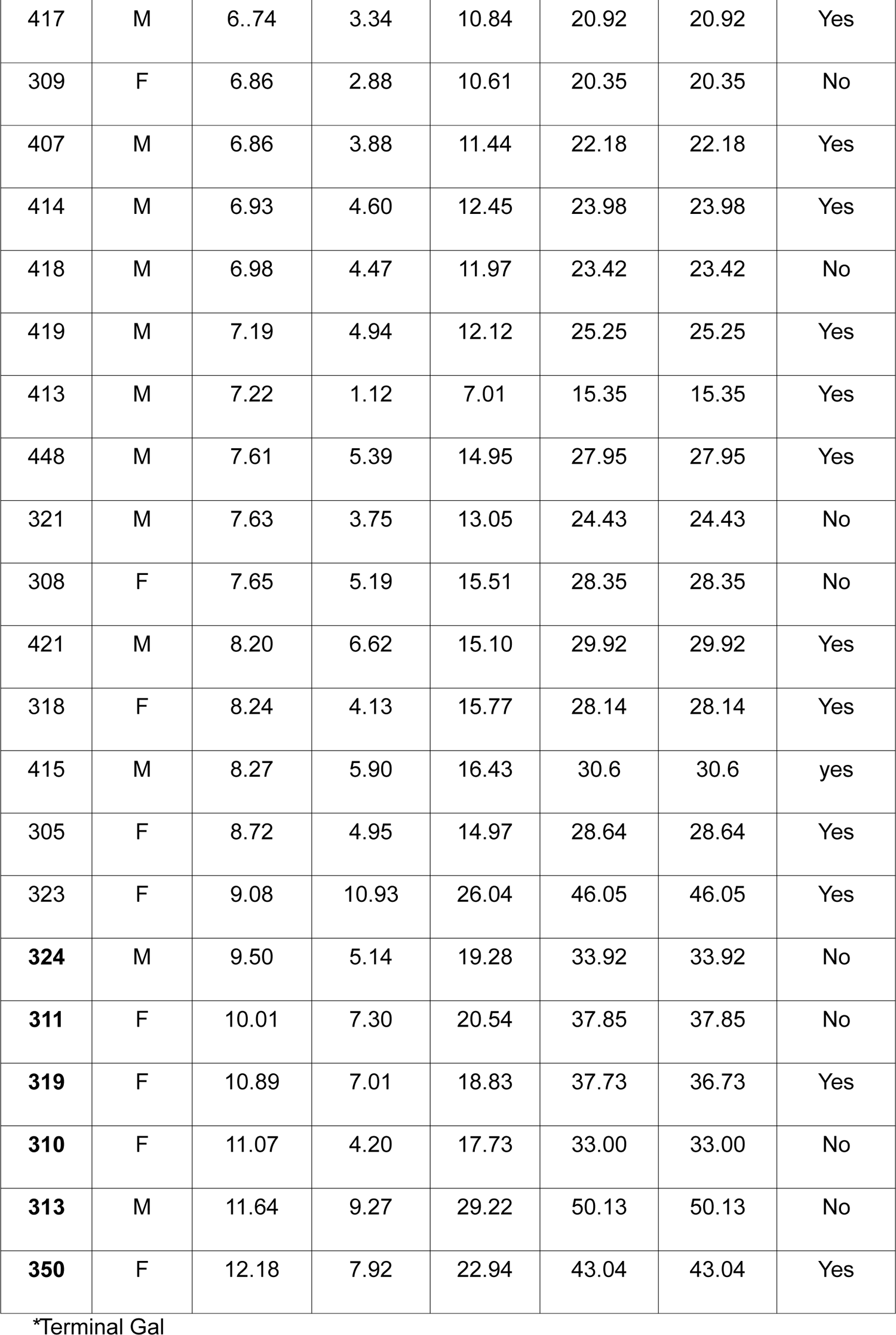
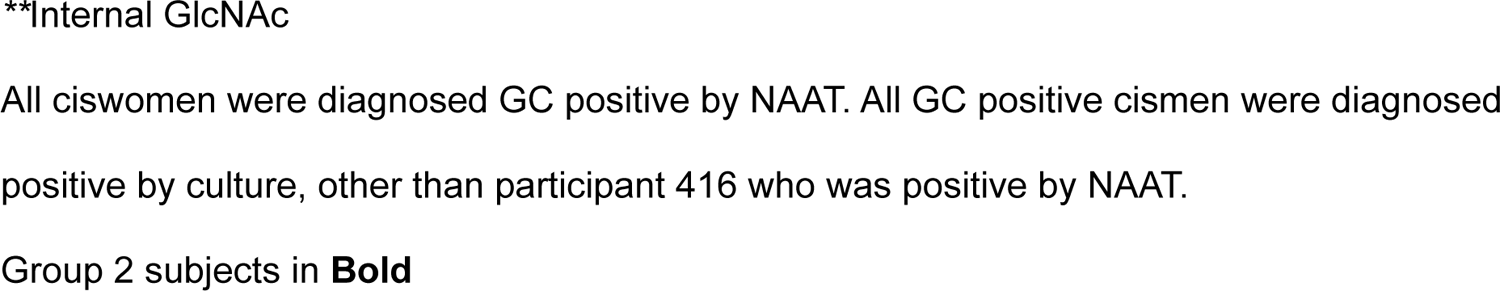
Concentrations of the Three Internal LOS IgG Specificities (ng/mL) Summed to that of IgG that bound the parent LOS.

### GC Culture & NAAT results

Seventy percent (26/37) of all GC-Contacts were infected, 12 of the 17 ciswomen (70%) and one cisman by Nucleic Acid Amplification Test and 13 cismen by culture (see Table 1 for participant numbers). Six ciswomen met Amsel criteria for bacterial vaginosis (BV)[19].

### Sexual Behavior and Clinical Presentation

The sexual behavior and clinical presentation of the 37 participants is summarized in Table 2. Fifty-four percent of GC-Contacts (20/37) reported to clinic within six days of exposure (Early Reporters), and 46% (17/37) waited seven days or longer to report (Delayed Reporters). Only vaginal pH and clinical BV discriminated Early Reporter ciswomen from Delayed Reporter ciswomen (Figure 2). Of the nine women who reported early and had a recorded vaginal pH, eight had a vaginal pH ≥ 5 and six had concurrent BV. Of the six women who delayed reporting and had a recorded vaginal pH, two had a vaginal pH ≥ 5 and none had concurrent BV. The difference in concurrent BV was significant (*p* = 0.03; Fisher’s Exact Test). Eighty-nine percent of cismen (8/9) who reported early, as well as 82% of cismen (9/11) who delayed reporting, complained of urethral discharge. GC infection status did not discriminate Eary verses Late Reporters in ciswomen; symptoms of BV likely prompted these ciswomen to present to clinical care sooner. However, urethral discharge in GC-infected cismen did differentiate Early verses Late Reporters.

**Figure 2.**
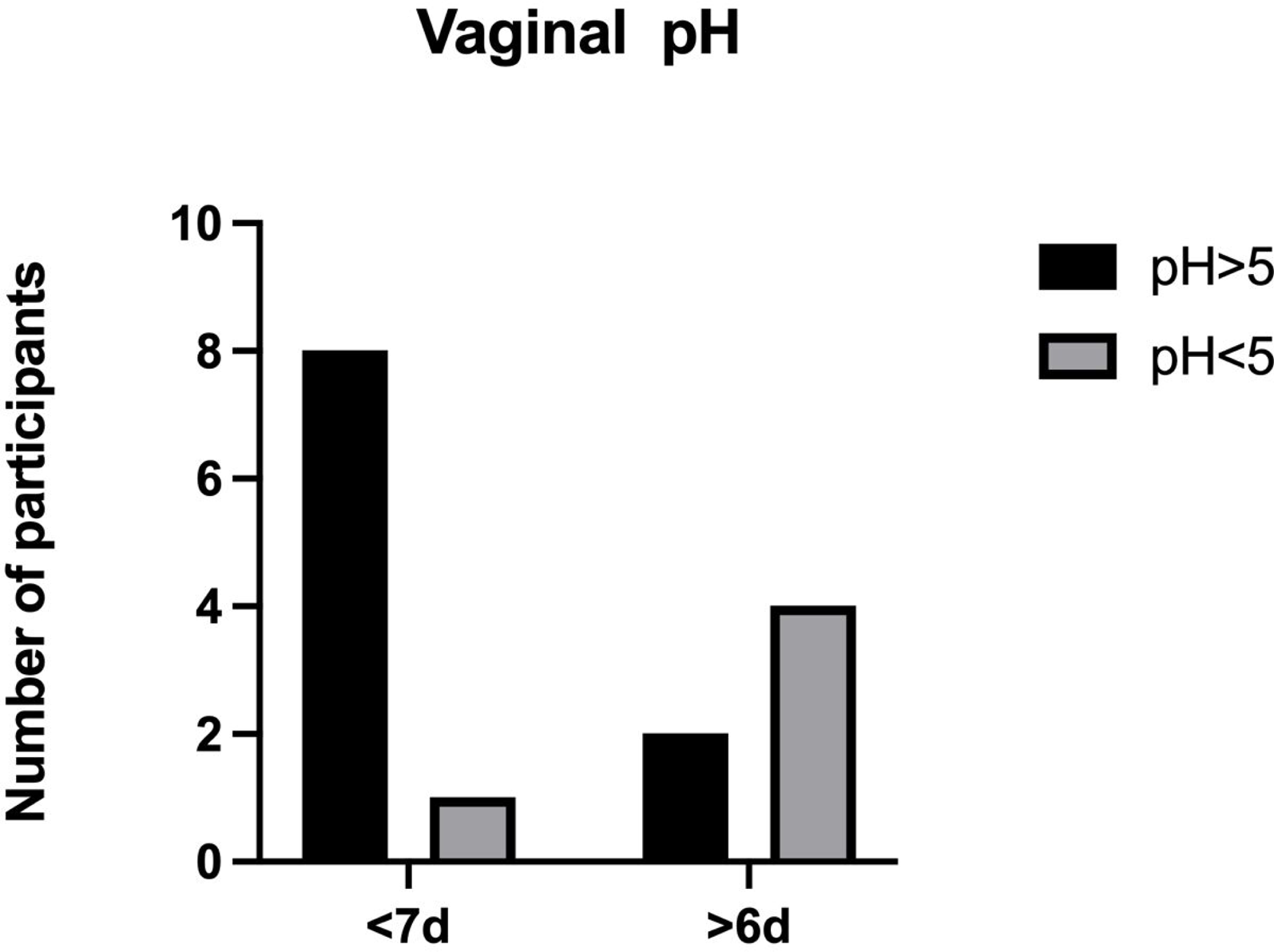
Vaginal pH of Early vs. Late Reporters.

**Table 2.** Sexual Behavior and Clinical Presentation of the 37 Participants.

|  |  |  |  |
| --- | --- | --- | --- |
|  |  | Cisgender Men<br>(n=20) | Cisgender Women<br>(n=17) |
| Sexual Behavior | Penis-vagina sex | 90% (18) | 100% (17) |
|  | Anal sex | 10% (2) | 6% (1) |
|  | Oral sex | 30% (6) | 41% (7) |
| Symptoms | Vaginal discharge and/or abdominal pain | N/A | 100% (17) |
|  | Urethral discharge | 85% (17) | N/A |
| Examination | Vaginal and/or cervical discharge | N/A | 47% (8) |
|  | Rectal discharge | N/A | 6% (1/17) |
|  | Bacterial Vaginosis (BV) | N/A | 35% (6/17) |
|  | Urethral discharge | 80% (16) | N/A |
| Diagnoses | Pelvic Inflammatory Disease (PID) | N/A | 18% (3) |
|  | PID with GC infection | N/A | 6% (1) |
|  | Cervical GC infection | N/A | 70% (12) |
|  | Penile urethral GC infection | 70% (14) | N/A |

### Multiplex Assay

Multiplexing the assay by combining multiple bead regions conjugated with different LOS did not result in interference between regions. The multiplexed and single-region assays gave nearly identical results.

The ability of the assay to accurately quantify IgG that bound only to different glycose antigens presented by the LOS molecule was confirmed by the fact that the concentrations of nLc4 IgG, Lc3 IgG and Lc2 IgG summed to the concentration of IgG that bound the parent 1291 LOS (Table 1).

### Serum α chain IgG Concentrations

The concentrations of nLc4 IgG ranged from 19.7% to 47.0% of the total 1291 IgG (mean: 22.8%); that of Lc3 IgG ranged from 7.3% to 24.2 % of the total 1291 IgG (mean: 17.9%).

During preliminary inspection of these data, we observed that the overall data distribution was not normal. Rather, these data appeared to cluster into two normal distributions. Determination of grouping as bimodal was conducted prior to knowledge of infection status and was the basis for our partitioning of Ab concentrations into Group 1 and Group 2.

The distribution of nLc4 and Lc3 IgG concentrations is shown in Figure 3. Thirty-one of the participants had nLc4 IgG concentrations < 9 ng/mL (mean: 6.22, SD. 1.58; Group 1), whereas six had >10 ng/mL (mean: 11.2, s.d. 0.818; Group 2). Several participants’ data arranged along the border of these 2 groups. They were partitioned, before statistical analysis of infection status and clinical presentation, according to standard deviations (s.d.) around the respective nLc4 IgG group means. The 9.08 ng/mL nLc4 concentration (Table 1 – participant 323) was within 2 s.d. of the Group 1 mean, and that participant was added to Group 1. Similarly, the 9.50 ng/mL (Table 1 – participant 324) concentration was 2 s.d. below the Group 2 mean and that participant was added to Group 2.

**Figure 3.**
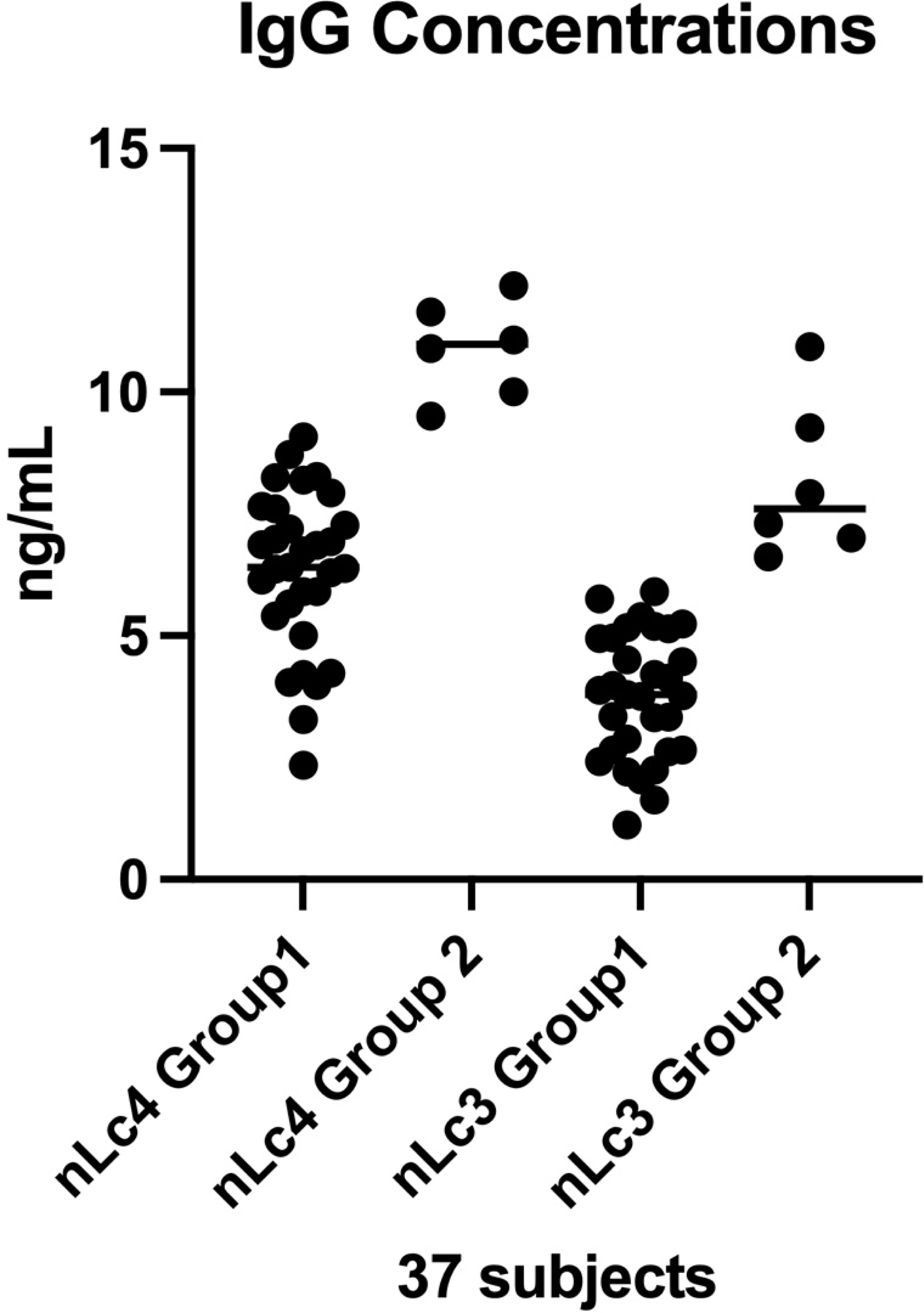
Distribution of serum nLc4 and Lc3 IgG concentrations among the 37 participants. Thirty-one of the participants had serum nLc4 IgG concentrations that were normally distributed around a mean of 6.33 ng/mL (SD: 1.65) and with a range of 2.34 to 9.08 ng/mL (Group 1); the other six participants (Group 2) had serum concentrations of nLc4 IgG that ranged from 9.50 to 12.18 ng/mL (mean:10.9, SD: 1.00). Serum concentrations of Lc3 IgG were similarly grouped: 31 participants had serum concentrations that were normally distributed around a mean of 3.76 ng/mL (SD: 1.28) with a range of 1.12 to 5.90 ng/mL (Group 1). The six Group 2 participants had serum concentrations that ranged from 6.62 to 10.9 ng/mL (mean:8.18, SD: 1.64). Four of the Lc3 IgG Group 2 participants, three cisgender females and one cisgender male, were also in the nLc4 IgG Group 2

Serum concentrations of Lc3 IgG similarly fell into two groups (Figure 3). Three ciswomen and one cisman in the Lc3 IgG Group 2 were also in the nLc4 IgG Group 2 The initial data inspection led us to question whether Group 2 differed from Group 1 in their response to GC exposure. The six Group 2 participants, four ciswomen and two cismen, differed from the Group 1 participants (Table 3). Group 2 participants were less likely to be infected (*p* = 0.05; Fisher’s exact test) and were mostly without abdominal pain or urethral or cervicovaginal discharge on exam (*p* = 0.03; Fisher’s exact test). They also were less likely to report a past gonococcal infection. Neither of the two cismen in Group 2, who had serum nLc4 IgG concentrations of 9.50 (Table 1 – participant 324) and 11.64 ng/mL (Table 1 – participant 313) were infected, though both complained of urethral discharge, and one was found to have discharge on exam. None of the four Group 2 women who had serum nLc4 IgG concentrations of 10.1-12.18 ng/mL complained of abdominal pain, and only one, who had BV, had vaginal discharge on exam. Both ciswomen who were infected in Group 2 had BV.

**Table 3.** Comparison of nLc4 IgG Group 1 and Group 2 Participants.

| Characteristic | nLc4 IgG Group 1 (*N* = 31) | nLc4 IgG Group 2 (*N* = 6) |  |
| --- | --- | --- | --- |
| Past gonococcal infection | 20/31 (65%) | 2/6 (33.3%) | *p* = 0.20 |
| Urethral or cervicovaginal discharge or abdominal pain | 25/31 (80.6%) | 2/6 (33.3%) | *p* = 0.03 |
| Infected | 24/31 (77%) | 2/6 (33.3%) | *p* = 0.05 |

### nLc4 and nLc3 concentration did not differ between cismen and ciswomen

nLc4 IgG concentrations in the sera of ciswomen did not differ from those in the sera of cismen. The mean nLc4 IgG of uninfected ciswomen was 7.9 ng/mL and of uninfected cismen was 7.5 ng/mL; The mean nLc4 IgG of infected ciswomen was 6.7 ng/mL and of infected cismen was 6.8 ng/mL. Lc3 IgG concentrations also did not differ between ciswomen and cismen.

### Gonococcal infections did not increase neo-antibody production against LOS

Infected Group 1 participants who presented more than six days after exposure would be expected to have made new antibodies in response to the infection[3], but concentrations of nLc4 and Lc3 IgG in their sera did not differ from those in the sera of Group 1 participants who reported within six days (means 6.27 and 6.38, respectively, for nLc4 IgG, and 4.37 and 4.07, respectively, for Lc3 IgG).

### Effect of prior gonococcal infections on nLc4 and Lc3 IgG concentrations

The was no evidence of a strong immunologic amnestic response to gonococcal infection (Figure 4). Twenty Group 1 participants (65%) had had 1-3 previous gonococcal infections. Those with past GC infections trend toward higher nLc4 and Lc3 IgG concentrations (Figure 4 and Supplemental Figure 1); although this is not significant. Given the small sample size, Group 2 Ab concentrations by past GC infection are shown in the supplemental Figure 1.

**Figure 4.**
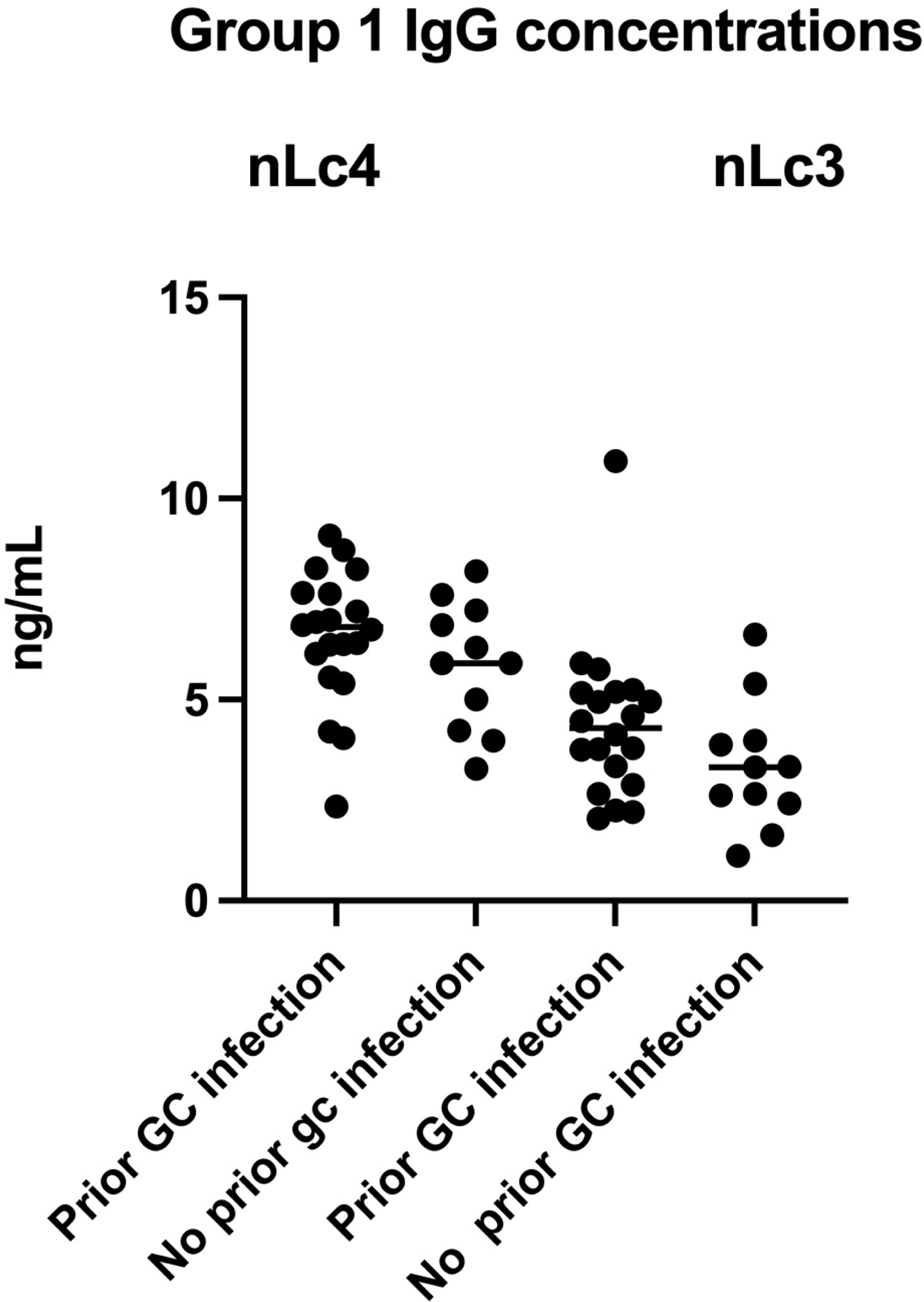
Effect of past GC infections - Group 1 Participants. Since Delayed Reporting was not associated with neo-antibody production, Early and Delayed Reporters are combined. There are no statistically significant differences in Lc3 or nLc4 IgG concentrations between those who had or had not had one or more previous GC infections. For participants who reported one or more prior GC infections, the mean nLc4 IgG concentration was 6.56 ng/mL (SD: 1.66); for those who had not had a prior infection, the mean was 5.66 ng/mL (SD: 1.59). For Lc3 IgG the corresponding values were 4.40 ng/mL (SD: 1.95) and 3.36 ng/mL (SD: 1.57), respectively.

## Discussion

The multiplexed fluorescent microsphere immunoassay developed by Luminex and provided by Bio-Rad has been widely used to simultaneously quantify antibodies binding to diverse antigens presented by proteins and has been adapted to simultaneously quantify antibodies that bind the several pneumococcal capsular polysaccharides in the sera of vaccinates[20]. However, it has not previously been adapted for the quantification of antibodies that bind the several antigens presented by a glycolipid such as LOS. The key adaptation was the use of deoxycholate to disperse LOS micelles[17] and permit the binding of individual LOS molecules to the fluorescent beads. Coupling LOS to the beads through the primary amino groups of their basal PEAs enables the oligosaccharide moieties to assume the minimal energy conformations that they would on the surface of a bacterium.

The assay was remarkably accurate. It quantified four IgG specificities: those that bound a native LOS with αn nLc4 α chain, IgG that bound only the non-reducing terminal galactose of that α chain (nLc4 IgG), IgG that bound only the internal glucosamine of that α chain (Lc3 IgG) and IgG that bound LOS with a truncated α chain that terminated at the internal β-Lactose (Lc2 IgG) (Figure 1). The first encompassed all the molecule’s antigens; the last, only those found in the basal portion of the molecule. Adding the concentrations of nLc4 and Lc3 IgG to that of Lc2 IgG in a participant’s serum exactly equaled the concentration of IgG that bound the native LOS molecule. Theoretically, it would be possible to inventory all the specificities of LOS IgG in any serum. This would but require an extended library of LOS mutants. One inherent limitation of this method is the possible alteration of the LOS epitopes by PEA binding, although binding to the PEA withing the lipid moiety should be less likely to affect these structures. It is possible that binding occurred through the PEA withing the carbohydrate structure. In future iterations of this assay, it would be pertinent to use a series of 1291 mutants that also have a *lptA* deletion to avoid this possibility[21].

We focused on nLc4 IgG and Lc3 IgG, because gonococci within PMNs in the urethral exudates of infected men make LOS with nLc4 α chains[8], nLc4 α chains are required for gonococcal invasion of epithelial cells *in vitro*[22,23], and IgG that bind nLc4 and Lc3 antigens have different functional activities against *N. meningitidis*[9]. We did not attempt to measure IgG specific for GalNAc-nLc4, an LOS α chain made by *N. gonorrhoeae* (Figure 1)[2], but not made by other *Neisseria*. To do so would have required additional mutants that were not available, and our objective was to develop and validate the assay. Given that the LOS biosynthetic genes are now known[10], making the necessary mutants would be straight-forward.

LOS-based vaccines to prevent gonococcal infections currently are in development, including one based on extended β chain LOS antigens[24]. This has been spurred by a reduced risk of gonococcal infections after puberty in young adults who were vaccinated with meningococcal outer membrane vesicle (OMV) vaccines as children[25]. OMV vaccines contain 6-9% LOS by weight. *N. meningitidis* OMV vaccine strains make LOS with nLc4 α chains.

Understanding the immunogenicity and effectiveness of these vaccines in humans will require an assay that detects antibodies to the different LOS antigens involved. Without such an assay, results of vaccine trials will be incomplete and may well be misleading, as the predominant response may be to an LOS antigen that does not induce protective antibodies[9]. The multiplexed LOS fluorescent microsphere immunoassay described herein fulfills this need. It can be readily expanded to quantify IgG specific for additional and different LOS antigens, as needed, by developing the needed LOS mutants.

The BCHD clients who participated in the larger study of which the present study is derived were typical of a large inner city sexually transmitted infection (STI) core: with many sexual partners from an early age and multiple previous STIs, including gonococcal infections[26,27]. Given the sexual risk profile of this core population, we expected higher concentrations of LOS IgG antibodies than we found. Only 22 % of participants had >30 ng/mL of IgG that bound 1291 LOS, and 16% had nLc4 IgG concentrations that fell into Group 2. We also were surprised by the negligible effect of prior gonococcal infections on serum concentrations of LOS IgG and the lack of a robust neo-antibody response to the current infection.

Hicks, *et al.*, previously had shown that acute gonorrhea resulted in new serum LOS IgG, detected by western blot analysis, in 6/13 men who had a first GC infection, and a marked increase in at least one who had preexisting LOS IgG[28]. The LOS antigen(s) that induced the IgG was not identified, but there were multiple bands in the LOS regions of the western blots. It may well be that the new IgG produced in response to those infections were specific for the GalNAc-nLc4 α chain or extended β chain antigens[2], which were not queried in our assay.

In contrast to the findings of Hicks, *et al.*, Hedges, *et al.,* using enzyme-linked immunosorbent assays, reported that, “in general, the local and systemic immune responses to [formalin-fixed] gonococci were extremely modest” and that prior gonococcal infections did not alter the response[29]. Hedges’ findings are consistent with what we found in the present study. It makes biologic sense that responses to foreign antigens by the organized lymphoid tissues of the uterine cervix and male urethra would be muted, as both are repeatedly exposed to allogeneic molecules, and the cervix must welcome allogeneic sperm. Hicks, *et al.*, also reported that asymptomatic pharyngeal colonization with *N. meningitidis* induced IgG that bound LOS and protein antigens in western blots[28]. Since most meningococcal strains make LOS with nLc4 α chains, this is a very plausible explanation for the presence of nLc4 α chain IgG in the sera of our participants. It also implies that the responses of the organized lymphoid tissues of the oral pharynx are not as restricted as those of the cervix and urethra.

IgG in endocervical and cervicovaginal secretions originates in serum; the mucosal associated lymphoid tissue of the female reproductive tract supplies only low levels of IgA[30]. After vaccination, serum HPV IgG is rapidly transudated into cervicovaginal secretions[31]. The serum concentrations of LOS IgG that we found would accurately reflect concentrations in cervicovaginal secretions, and the assay described here can be used to measure the serum response to any future LOS-based vaccines with confidence that the quantities reflect those in cervicovaginal secretions. The relationship between serum IgG concentrations and those in seminal fluids is not understood.

The Group 2 participants were interesting. They had higher concentrations of LOS IgG than Group 1 participants and were less likely to be infected during exposure. It will require a much larger study to establish an accurate IgG concentration boundary between Groups 1 and 2 and determine the true effect of serum nLc4 IgG concentrations ≥ 9.5 ng/mL on the risk of acquiring a gonococcal infection during exposure at different anatomic sites. The molecular environments of the cervix and vagina are far more complex than that of the male urethra[32], and it is unlikely that nLc4 IgG, or any LOS antibody, will have the same effect in men as in women.

In summary we have developed a multiplex assay that accurately quantifies serum concentrations of IgG that bind three different glycose antigens in the *Neisseria* LOS nLc4 α chain. The assay can be expanded to encompass additional LOS antigens with the development of the necessary LOS mutants, and it can be used to inventory LOS IgG in a serum, determine the effects of different LOS IgG specificities on susceptibility and resistance to gonococcal infections, and analyze responses to LOS-based vaccines that are in development. We have used it in a preliminary study to inform larger studies of the effects of gonococcal LOS IgG on susceptibility and resistance to infection.

---

**Supplemental Figure 1.** Group 2 anti-nLc4 and anti-Lc3 IgG Concentrations by prior gonococcal infection status.

## Footnotes

1 Potential Conflicts of Interest: Stephanie E. McLaughlin has no potential conflicts of interest to declare. Zhijie Yang has no potential conflicts of interest to decline, and J. McLeod Griffiss has no potential conflicts of interest to declare.

2 This work was supported, in whole or in part, by National Institutes of Health Grants AI 053728 and AI 065605 (J.Mcl.G, P.I.) administered by the Northern California Institute for Research and Education and with resources of the Veterans Affairs Medical Center (San Francisco, CA). This is publication 113 from the Center for Immunochemistry.

3 These data were presented at the 23^rd^ International Pathogenic Neisseria Conference (Boston MA, USA 2023).

## References

1. Schmidt KA, Schneider H, Lindstrom JA, et al. Experimental Gonococcal Urethritis and Reinfection with Homologous Gonococci in Male Volunteers. Sex Transm Dis. 2001; 28(10):555–564.

2. Schneider H, Griffiss JM, Boslego JW, Hitchcock PJ, Zahos KM, Apicella MA. Expression of para-globoside-like lipooligosaccharides may be a necessary component of gonococcal pathogenesis in men. J Exp Medicine. 1991; 174(6):1601–1605.

3. Barrier BF, Gargiulo AR, Schust DJ. Reproductive Immunology and Its Disorders. Chapter 13 In: Yen & Jaffe’s Reproductive Endocrinology. Strauss JF, Barbieri RL, editors. 6th ed. 2009. p. 299–323.

4. Griffiss JM, Schneider H, Mandrell RE, et al. Lipooligosaccharides: The Principal Glycolipids of the Neisserial Outer Membrane. Clin Infect Dis. 1988; 10(Supplement 2): S287–S295.

5. Griffiss JM. The Role of Bacterial Lipooligosaccharides in the Pathogenesis of Human Disease. Trends Glycosci Glycotechnol. 2010; 7(38):461.

6. Griffiss JMcL, Schneider H. Endotoxin in Health and Disease. The chemistry and biology of lipooli-gosaccharides: The endotoxins of bacteria of the respiratory and genital mucosae. New York: Marcel Dekker; p. 174–194.

7. Mandrell RE, Griffiss JM, Macher BA. Lipooligosaccharides (LOS) of Neisseria gonorrhoeae and Neisseria meningitidis have components that are immunochemically similar to precursors of human blood group antigens. Carbohydrate sequence specificity of the mouse monoclonal antibodies that recognize crossreacting antigens on LOS and human erythrocytes. J Exp Medicine. 1988; 168(1):107–126.

8. McLaughlin SE, Cheng H, Ghanem KG, et al. Urethral Exudates of Men with Neisseria gonorrhoeae Infections Select a Restricted Lipooligosaccharide Phenotype During Transmission. J Infect Dis. 2012; 206(8):1227–1232.

9. Cheng H, Yang Z, Estabrook MM, et al. Human Lipooligosaccharide IGG That Prevents Endemic Meningococcal Disease Recognizes an Internal Lacto-N-neotetraose Structure. J Biol Chem. 2011; 286(51):43622–43633.

10. Yang QL, Gotschlich EC. Variation of gonococcal lipooligosaccharide structure is due to alterations in poly-G tracts in lgt genes encoding glycosyl transferases. J Exp Medicine. 1996; 183(1):323– 327.

11. Griffiss JM, O’Brien JP, Yamasaki R, Williams GD, Rice PA, Schneider H. Physical heterogeneity of neisserial lipooligosaccharides reflects oligosaccharides that differ in apparent molecular weight, chemical composition, and antigenic expression. Infect Immun. 1987; 55(8):1792–1800.

12. Yamasaki R, Schneider H, Griffiss JMcL, Mandrell R. Epitope expression of gonococcal lipooligosaccharide (LOS). Importance of the lipoidal moiety for expression of an epitope that exists in the oligosaccharide moiety of LOS. Mol Immunol. 1988; 25(8):799–809.

13. Schmiel DH, Moran EE, Keiser PB, Brandt BL, Zollinger WD. Importance of Antibodies to Lipopolysaccharide in Natural and Vaccine-Induced Serum Bactericidal Activity against Neisseria meningitidis Group B. Infect Immun. 2011; 79(10):4146–4156.

14. Yamasaki R, Yabe U, Kataoka C, Takeda U, Asuka S. The Oligosaccharide of Gonococcal Lipooligosaccharide Contains Several Epitopes That Are Recognized by Human Antibodies▿. Infect Immun. 2010; 78(7):3247–3257.

15. John CM, Griffiss JM, Apicella MA, Mandrell RE, Gibson BW. The structural basis for pyocin resistance in Neisseria gonorrhoeae lipooligosaccharides. J Biol Chem. 1991; 266(29):19303–19311.

16. McLaughlin SE, Ghanem KG, Zenilman JM, Griffiss JM. Risk of Gonococcal Infection During Vaginal Exposure is Associated with High Vaginal pH and Active Menstruation. Sex Transm Dis. 2019; 46(2):86–90.

17. Bertram MA, Griffiss JM, Broud DD. Response to Antigenic Determinants of Neisseria Meningitidis Lipopolysaccharide Investigated with A New Radioactive Antigen-Binding Assay. J Immunol. 1976; 116(3):842–846.

18. Bio-Rad. Multiplex Immunoassays. [Internet]. [cited 2024 Aug 17]. Available from: www.bio-rad.com/en-us/applicationtechnologies/multiplex-immunoassays?ID=LUSM0E8UU

19. Amsel R, Totten PA, Spiegel CA, Chen KCS, Eschenbach D, Holmes KK. Nonspecific vaginitis Diagnostic criteria and microbial and epidemiologic associations. Am J Medicine. 1983; 74(1):14–22.

20. Pickering JW, Martins TB, Greer RW, et al. A Multiplexed Fluorescent Microsphere Immunoassay for Antibodies to Pneumococcal Capsular Polysaccharides. Am J Clin Pathol. 2002; 117(4):589–596.

21. Lewis LA, Choudhury B, Balthazar JT, et al. Phosphoethanolamine Substitution of Lipid A and Resistance of Neisseria gonorrhoeae to Cationic Antimicrobial Peptides and Complement-Mediated Killing by Normal Human Serum▿. Infect Immun. 2009; 77(3):1112–1120.

22. Wang J. Neisseria gonorrhoeae must express the paraglobosyl LOS in order to invade human genitourinary epithelial cells. In: WD Z, CE F, CD D, editors. Abstracts of the Tenth International Pathogenic Neisseria Conference. Baltimore, Maryland; 1996.

23. Song W, Ma L, Chen R, Stein DC. Role of Lipooligosaccharide in Opa-Independent Invasion of Neisseria gonorrhoeae into Human Epithelial Cells. J Exp Medicine. 2000; 191(6):949–960.

24. Gulati S, Shaughnessy J, Ram S, Rice PA. Targeting Lipooligosaccharide (LOS) for a Gonococcal Vaccine. Front Immunol. 2019; 10:321.

25. Whelan J, Kløvstad H, Haugen IL, van Holle MR-DR B, Storsaeter J. Ecologic Study of Meningococcal B Vaccine and Neisseria gonorrhoeae Infection, Norway. Emerg Infect Dis. 2016; 22(6):1137– 1139.

26. Aral SO. Behavioral Aspects of Sexually Transmitted Diseases. Sex Transm Dis. 2000; 27(6):327– 328.

27. Thomas JC, Tucker MJ. The Development and Use Of The Concept Of A Sexually Transmitted Disease Core. J Infect Dis. 1996; 174(Supplement_2): S134–S143.

28. Hicks CB, Boslego JW, Brandt B. Evidence of Serum Antibodies to Neisseria gonorrhoeae Before Gonococcal Infection. J Infect Dis. 1987; 155(6):1276–1281.

29. Hedges SR, Mayo MS, Mestecky J, Hook EW, Russell MW. Limited Local and Systemic Antibody Responses to Neisseria gonorrhoeae during Uncomplicated Genital Infections. Infect Immun. 1999; 67(8):3937–3946.

30. Bard E, Riethmuller D, Biichlé S, et al. Validation of a Highly Sensitive Immunoenzymatic Assay to Establish the Origin of Immunoglobulins in Female Genital Secretions. J Immunoass Immunochem. 2002; 23(2):145–162.

31. Schwarz TF, Kocken M, Petäjä T, et al. Correlation between levels of human papillomavirus (HPV)-16 and 18 antibodies in serum and cervicovaginal secretions in girls and women vaccinated with the HPV-16/18 AS04-adjuvanted vaccine. Hum Vaccines. 2010; 6(12):1054–1061.

32. Marrazzo J. Doxycycline Postexposure Prophylaxis for STIs in Women — Uncertain Benefit, Urgent Need. N Engl J Med. 2023; 389(25):2389–2390.

